# Intracranial EEG reveals bihemispheric parietal and extra-parietal brain networks supporting mental arithmetic

**DOI:** 10.1101/542142

**Authors:** Gray S. Umbach, Michael D. Rugg, Gregory A. Worrell, Michael R. Sperling, Robert E. Gross, Barbara C. Jobst, Josh Jacobs, Kareem Zaghloul, Joel Stein, Kathryn Davis, Bradley C. Lega

## Abstract

Mathematical reasoning is central to everyday life. Lesion data and functional MRI studies suggest that even simple arithmetic involves the coordination of multiple spatially diverse brain regions, but to date, math processing has not been well characterized using direct brain recordings, especially outside of the parietal cortex. To address this, we utilized an unparalleled data set of 310 subjects implanted with intracranial electrodes to investigate the spatial and temporal dynamics of arithmetic reasoning. Our data support the importance of regions previously implicated in numerosity such as the superior parietal lobule and intraparietal sulcus. However, we also identify contributions to arithmetic processing from regions such as the entorhinal cortex and temporal pole. Using the excellent bihemispheric coverage afforded by our data set and the precise temporal resolution of intracranial recordings, we characterize in detail the spatial and temporal characteristics of an arithmetic processing network, quantifying subtle hemispheric differences for selected regions. We also examine how activity in these regions predicts arithmetic ability and look for gender differences in the pattern of network activation. Our findings further define the complex network of regions involved in human arithmetical reasoning.

## 1. Introduction

Mathematical processing is a central feature of human cognition, present across societies and linguistic systems. Its neural bases have been characterized through numerous noninvasive imaging and lesion studies. These data have suggested that mathematical manipulation requires complex interaction among bihemispheric brain regions, including understanding of value associated with numerical symbols, semantic memory of mathematical facts, the manipulation of numerical quantities using an “internal number line,” spatial attentional resources, and the manipulation of numerical quantities in working memory (Dehaene 1992; Dehaene et al. 2003; Daitch et al. 2016). It is thought that mathematical reasoning depends and even builds upon brain regions involved other cognitive processes such as non-verbal quantity manipulation and finger counting and as such parietal regions are most active during arithmetical cognition (Berteletti and Booth 2016; Nieder and Dehaene 2009). Central to this proposed parietal circuitry is the horizontal intraparietal sulcus (HIPS), a region that has exhibited reliable activation for tasks requiring specific manipulation of quantities, estimation, and equation solving (Dehaene 1992; Dehaene et al. 1999; Piazza et al. 2004; Dastjerdi et al. 2013; Amalric and Dehaene 2016; Daitch et al. 2016). These data have led some investigators to propose that the HIPS is specifically and exclusively active during mathematical cognition, while dissenting scientists believe it is more generically activated by quantity estimation and manipulation including spatial and other quantities. The HIPS is necessary but not sufficient for mathematical reasoning, according to the “three parietal circuits” model proposed by Dehaene (Dehaene et al. 2003). Along with the “internal number line” anatomically centered in the HIPS, mathematical tasks also recruit more dorsal parietal regions (superior parietal lobule, precuneus) which are thought to provide an orienting signal for math and spatial tasks as a specific functional domain within wider dorsal attention networks. The SPL however may also support ongoing mathematical processing apart from this initial attention function, as lesions in the SPL can elicit deficits on spatial and numerical bisection tasks (Zorzi et al. 2002). The final contributor to this three-circuit model is the angular gyrus, which supports semantic comprehension and representation of the numerical values associated with mathematical symbols (Grabner et al. 2009). A dissociation between angular gyrus and HIPS/SPL activation has been observed experimentally using paradigms that distinguish between mathematical fact retrieval (simple addition or multiplication of likely memorized equations) and more conventional numerical processing (such as estimation of quantity) (Dehaene et al. 1999; Grabner et al. 2009). Associated with these parietal circuit is an occipitotemporal region thought to preferentially activate during numerical item presentation (the putative number form area, NFA) (Shum et al., 2013).

The three parietal regions plus NFA map partially onto a proposed cognitive “division of labor” during mathematical processing, by which visual input must be decoded (interpretation of Arabic numeral code NFA), semantic information of mathematical statements (especially memorized expressions) must be extracted/interpreted (semantic code angular gyrus/semantic language regions), and quantities must be represented along an internal number line and be evaluated and manipulated (HIPS). While this combined triple code/parietal circuits model incorporates the most active mathematical processing areas, it is highly likely that other brain regions may contribute as well. Some examples include the prefrontal cortex, which has been observed to house neurons tuned to specific numerosities in primates (Nieder 2012). For more complex arithmetical manipulations, this region may also play a role in working memory maintenance of intermediate values during equation solving (Curtis and D’Esposito 2003). Likewise, at least during development, regions of the temporal lobe may participate in semantic representations of mathematical symbols or numbers (Menon 2016), and noninvasive studies have observed activation in other regions such as the insula (Simon et al. 2002), implying some correlation of activity in rhinal and limbic networks with arithmetical processing, though a specific role has not been defined for these within a math processing network. Importantly, the involvement of these and other extra-parietal regions in arithmetic has not been characterized using direct brain recordings (Daitch et al. 2016).

While there is no evidence for difference in math aptitude between genders (Hyde et al. 2008), a noninvasive imaging study using an arithmetic task demonstrated greater activation of the right posterior parietal cortex structures in males compared to females (Keller and Menon 2009), suggesting potential divergence of problem-solving strategy. We know of no iEEG study that has explored differences in arithmetic problem-solving strategy further.

Existing human iEEG studies of arithmetical processing have been limited by inclusion of relatively few subjects, but these investigations have supported the existence of specifically numerically sensitive regions in the inferior temporal area and of the importance of the IPS to arithmetical processing (Daitch et al. 2016; Pinheiro-Chagas et al. 2018). However, several components of the proposed functional-neuroanatomic models have not been investigated, such as unilateral versus bilateral activation of these parietal regions. Further, the exclusive use of subdural grid electrodes in prior studies precludes sampling from subcortical locations and the possibility of disambiguating activity arising from midline parietal structures (namely the precuneus) from that generated in the dorsal SPL. Depth electrodes also aid in differentiating dorsal midline structures like the precuneus, expected to be active in arithmetic (Dehaene et al. 2003), from contributions of the posterior cingulate cortex, shown to be suppressed during arithmetic (Foster et al. 2012). Finally, the possible participation of brain regions in an arithmetic processing network outside of the ventral temporoparietal cortex has not been investigated using direct brain recordings, in spite of extensive functional imaging evidence indicating that other brain regions (especially the pre-frontal cortex and insula) exhibit mathematics related activation. Support for the importance of direct recordings to characterize arithmetical processing regions has come from a recent electrophysiology study indicating that the temporal dynamics of activation in some brain regions may render the activity undetectable to analyses based solely upon BOLD signal changes (Pinheiro-Chagas et al. 2018).

To address these gaps in the literature surrounding electrophysiological activity underlying arithmetical processing in humans, we analyzed an unparalleled data set of 310 subjects who performed an arithmetical task during monitoring with intracranial electrodes for seizure mapping. We identified a core arithmetical processing network by identifying brain locations that show significantly greater activity during arithmetical reasoning as compared to successful episodic memory encoding. The advantage of this comparison condition is that we are able to reduce the influence of nonspecific task-related activity in the brain (such as that associated with focusing of attention), although this paradigm may underestimate the contribution brain areas involved in retrieval of semantic facts (see discussion). In addition, our data set included over 50 subjects with parietal cortex depth electrodes inserted via the stereo EEG technique, allowing us to obtain signal directly from the IPS and to disambiguate the contributions to arithmetical processing of dorsal SPL locations from those of midline parietal areas (precuneus and posterior cingulate). Using direct brain recordings, we were able to explore in detail temporal dynamics within a core arithmetic processing network as well as how these temporal patterns differed across the hemispheres. The large number of participants allowed us to examine how activity within regions predicted math performance, and also to look for gender-related differences in processing patterns.

## 2. Methods

### 2.1 Participants

A total of 310 adult patients were consented to contribute to the study. Patients were recruited as part of a national research consortium for the investigation of memory processing (DARPA RAM program) as well as through iEEG studies conducted independently by this program at UT Southwestern and the University of Pennsylvania. Patient characteristics including handedness and hemispheric dominance were assessed by local clinical experts.

### 2.2 Behavioral task

Subjects performed the free recall task on a laptop computer. The laptop transmitted serial electrical pulses into the clinical system to allow for alignment of behavioral events to the corresponding iEEG data. Several variations of this task provided data for this analysis, with minor differences in the total number of memory items and categorical structure of the study list. However, the arithmetic component of these tasks was identical across all variants of the recall procedure. In all cases, subjects were presented with a series of words for later recall. All the words of a list (12-15 items) were capitalized and displayed at the center of the screen for 1600 ms. Each word was followed by an 400-600 ms inter-stimulus interval during which there was no stimulus on the screen. After the final word, patients immediately engaged in a series of arithmetic problems of the form A + B + C = ?? for a minimum duration of 20 seconds, with A, B, and C representing randomly selected integers ranging from one to nine. Participants entered their answer via the laptop’s keyboard. A new arithmetic problem populated the screen as soon as the previous answer was entered. Correct reposes took an average of 6.55 seconds, and subjects answered correctly 91.2% of the time, meaning roughly 3 arithmetic problems per 20 second period were displayed and solved. After completion of the final arithmetic problem, a tone signaled the beginning of the free recall period, lasting 45 seconds, during which patients were required to verbalize as many words from the most recent list as they could remember. The patients performed as many lists as they were able in a given testing session. The comparison we made to identify arithmetic-related processing networks was between successful arithmetic events (participant entered correct answer) and successful memory encoding events (encoding event of a word later recalled verbally). A schematic of the task is shown in Figure 1A. A single arithmetic consisted of the first 2 seconds during which an arithmetic problem was displayed. A single encoding event consisted of the first 2 seconds during which a single word was displayed during the encoding period of the task. No subject included in the analysis had an average response time to the arithmetic problems faster than 2200 ms, allowing for a 2 second time window to be used for the comparison between tasks.

**Figure 1:**
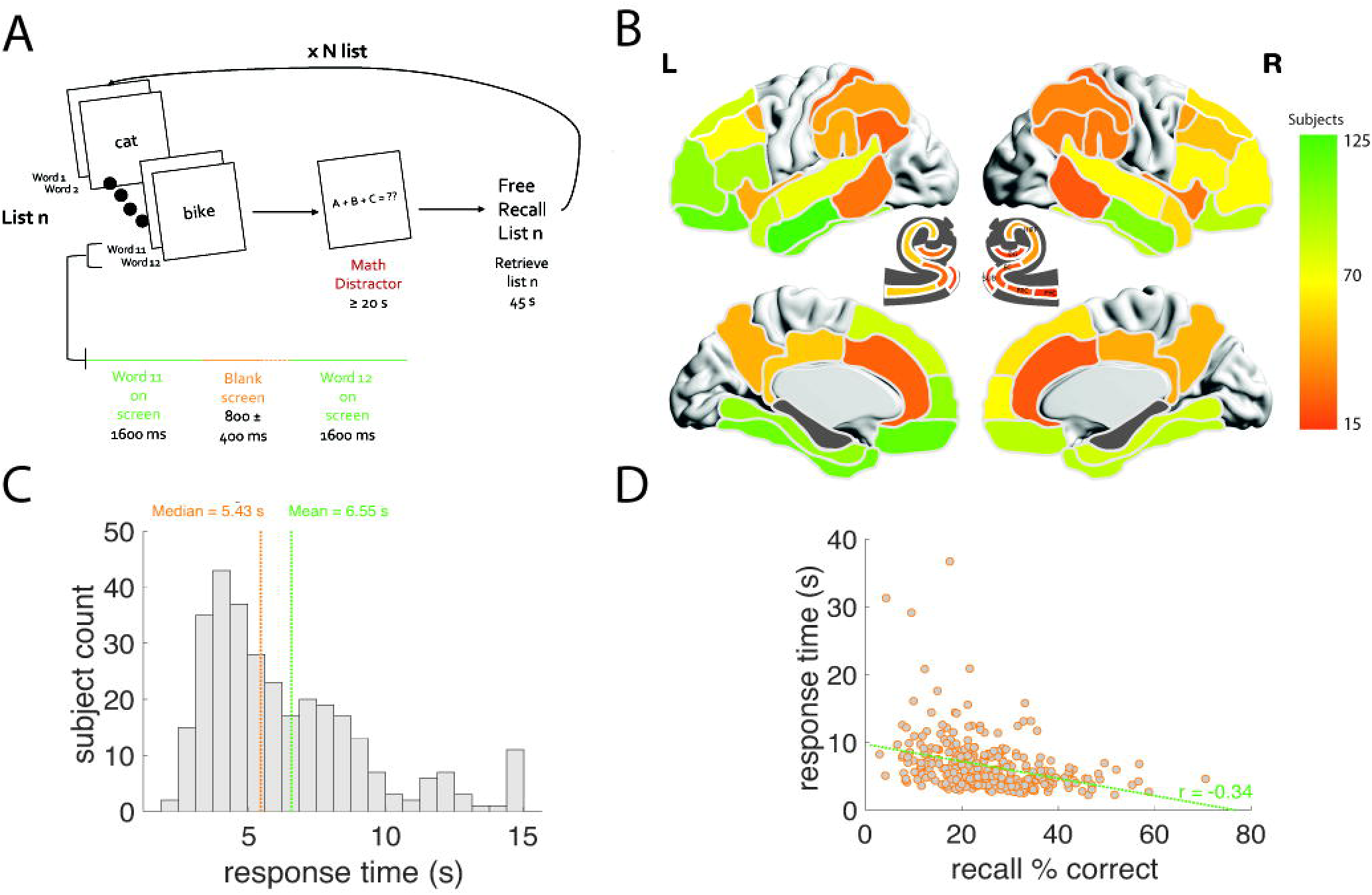
Task, Subject Count, and Behavioral Data. **A.** Schematic of free recall task. **B.** Subject count for each region included in analysis. We a priori excluded primary occipital and Rolandic areas from the analysis. **C.** Distribution of response times arithmetic problems with mean and median superimposed. **D.** Subject level correlation of performance during memory task as measured by percent of words correctly recalled with performance during the arithmetic task as measured by the average response time during the arithmetic problems.

### 2.3 Brain regions

Both surface intracranial electrodes (subdural grids and strips) and depth electrodes (principally stereo EEG probes) were included in the analysis. Electrode localization was accomplished by co-registering postoperative CT scans with a standard Talairach atlas. X, Y and Z locations in Talairach space were derived for all electrodes included. In addition, the location of all electrodes underwent manual expert neuroradiology review for sub localization within the mesial temporal lobe or subcortical parietal regions, including the IPS. Electrodes were excluded from the analysis if they were present in areas of seizure onset, exhibited excessive interictal artifact or excessive electrical noise. Erroneously localized electrodes (those for which X, Y, Z coordinates placed the electrodes far outside the brain surface for example) were excluded from the spatial analysis. Electrode locations were reviewed manually using BrainNet Viewer in Matlab (Xia et al. 2013). Electrode classification into brain regions for analysis was accomplished via a combination of expert neuroradiological review (anatomical description of an electric location collected at the time of initial implantation) and automated localization using Talairach coordinates, region labels, and Brodmann areas. Direct review was especially critical for all depth electrodes, when anatomical location can be challenging and requires a good understanding of three-dimensional anatomy within sulci. Direct review was especially important for electrodes located within the IPS, for which no automatic label existed. For subdural strip and grid electrodes, XYZ Talairach localizations were used to automatically assign electrodes to Broadmann areas and anatomical regions (e.g. “superior frontal gyrus,” “middle temporal gyrus.”) We eliminated any electrodes for which this localization was not in agreement, for example an electrode localized to Brodmann area 20 but labeled as belonging to the “superior temporal gyrus.” We used both of these anatomical and Broadmann localizations to classify the electrodes in order to more narrowly define regions (compared to using only gyral/sulcal anatomy or Broadmann areas in isolation). We made the a priori decision to subdivide the temporal lobe and temporo-occipital regions as follows: the temporal pole (defined as Broadman area 38), the superior temporal gyrus (electrodes labeled as both STG and Brodmann area 22), Heschl’s gyrus/primary auditory region (defined as STG and Brodmann 41 and 42), the inferior temporal region (analyzed continuously along the y-axis, described below, including Broadmann areas 20 and 37 but excluding electrodes localized to the fusiform gyrus which was analyzed as a separate region), and the anterior and posterior middle temporal gyrus (dividing electrodes localized to the MTG between BA 21 and 37). The supramarginal gyrus (BA 40), and the angular gyrus (BA 39) were analyzed separately. For the frontal lobe, we separately analyzed dorsomedial prefrontal cortex, dorsolateral prefrontal cortex, ventral lateral prefrontal cortex, the frontal pole, and the orbitofrontal cortex. These were defined as follows: DLPFC included middle frontal gyrus electrodes localized to BA 46 and 9, DMPFC included superior frontal gyrus electrodes localized to BA 9 and 8, VLPFC included inferior frontal gyrus electrodes localized to BA 44, 45, and 46 and BA 9 and 46 of the inferior frontal gyrus, the frontal pole included any frontal lobe electrodes in BA 10, and orbitofrontal cortex included any frontal lobe electrodes assigned to Broadman area 11, 12, or 47. We made the a priori decision to exclude any electrodes localized to the primary motor or sensory cortex, and to excluded electrodes localized to the supplemental motor area (“SMA” label) or BA 6. We also excluded occipital regions from the analysis due to lack of coverage in core calcarine areas. All electrodes from mesial temporal structures including the hippocampus, entorhinal cortex (EC), and parahippocampus were localized via expert review and analyzed separately. Surface electrodes localized to assigned to the IPS were manually reviewed. All electrodes assigned to either the anterior or posterior cingulate regions were depth electrodes, the localization of which was manually determined. We excluded any electrode in the “mid–cingulate” region, defined using the paracentral sulcus and marginal sulcus (ascending ramus of the cingulate sulcus) as points of demarcation, although in practice anterior cingulate electrodes were located adjacent to the genu while posterior cingulate electrodes were adjacent to the splenium. Using these careful processes, we were able to take advantage of extensive expert neuroradiology input in our electrode locations, ensuring that electrodes and parietal regions were accurately localized within areas exhibiting arithmetical related activity. Electrodes demonstrating non-electrophysiological signal noise were excluded as part of a standard preprocessing pipeline.

### 2.4 Intracranial EEG processing

Multiple clinical systems were used for iEEG recording, depending on site, including systems from Nihon-Kohden, Nicolet, Grass Telefactor, DeltaMed (Natus), and Bio-Logic. Sampling frequency ranged from 250 to 2000 Hz, depending on clinical needs. Analog pulses were sent from the experimental laptop to the clinical system, allowing for subsequent aligning of experimental events to the EEG signals. Morlet wavelets were used to decompose the EEG signals into 57 logarithmically spaced frequencies ranging from 1-128 Hz. 500 ms buffers surrounding events were used to avoid edge artifacts, and a 60 Hz notch filter was used to reduce signal noise. EEG signals were downsampled to 500 Hz to reduce computational burden. Matlab software was used for all analyses. We made the a priori decision to focus on oscillatory activity in the high gamma frequency range (high-frequency activity or HFA, 64-128Hz) based upon pre-existing data (Dastjerdi et al. 2013; Daitch et al. 2016).

### 2.5 Whole brain analysis

Our principal goal was to characterize an arithmetical processing network. For this we sought to identify brain regions with greater activity for arithmetic-related processing by directly comparing oscillatory power during arithmetic problem-solving power recorded during study events associated with correct later recall (successful encoding). Power was extracted across the time series (downsampled to 500 Hz after power extraction). These power values were logarithmically transformed to generate normally distributed power distributions. A two-sample, one-tailed t-test across trials for a single electrode was performed between the parametric power distributions derived from arithmetic events and power associated with successful encoding. This generated a time-frequency matrix of p-values for each experimental protocol of each electrode of each subject quantifying the significance of the difference in power between the two classes of trial (arithmetic vs. encoding). The Z values obtained from these p-values were averaged across all electrodes for each region and subject. We then averaged across the entire 2 second time series to get a single Z value per subject in each region. We then tested this distribution of Z values versus a null hypothesis of no difference between the two conditions (Z value of 0). We obtained our main metric of brain activity by limiting to this analysis to the high frequency band (64 – 128 Hz). We named this metric ‘high frequency activity’ (HFA). We FDR corrected the resulting P values from a one sample t-test (Q = 0.01) to identify regions exhibiting significantly greater arithmetical activity. Each subject contributing data to a given brain region contributed only a single value; the number of subjects contributing to each brain region included in the analysis are shown in Figure 1B. For the initial analysis shown in figure 2, we analyzed the hemispheres separately.

**Figure 2:**
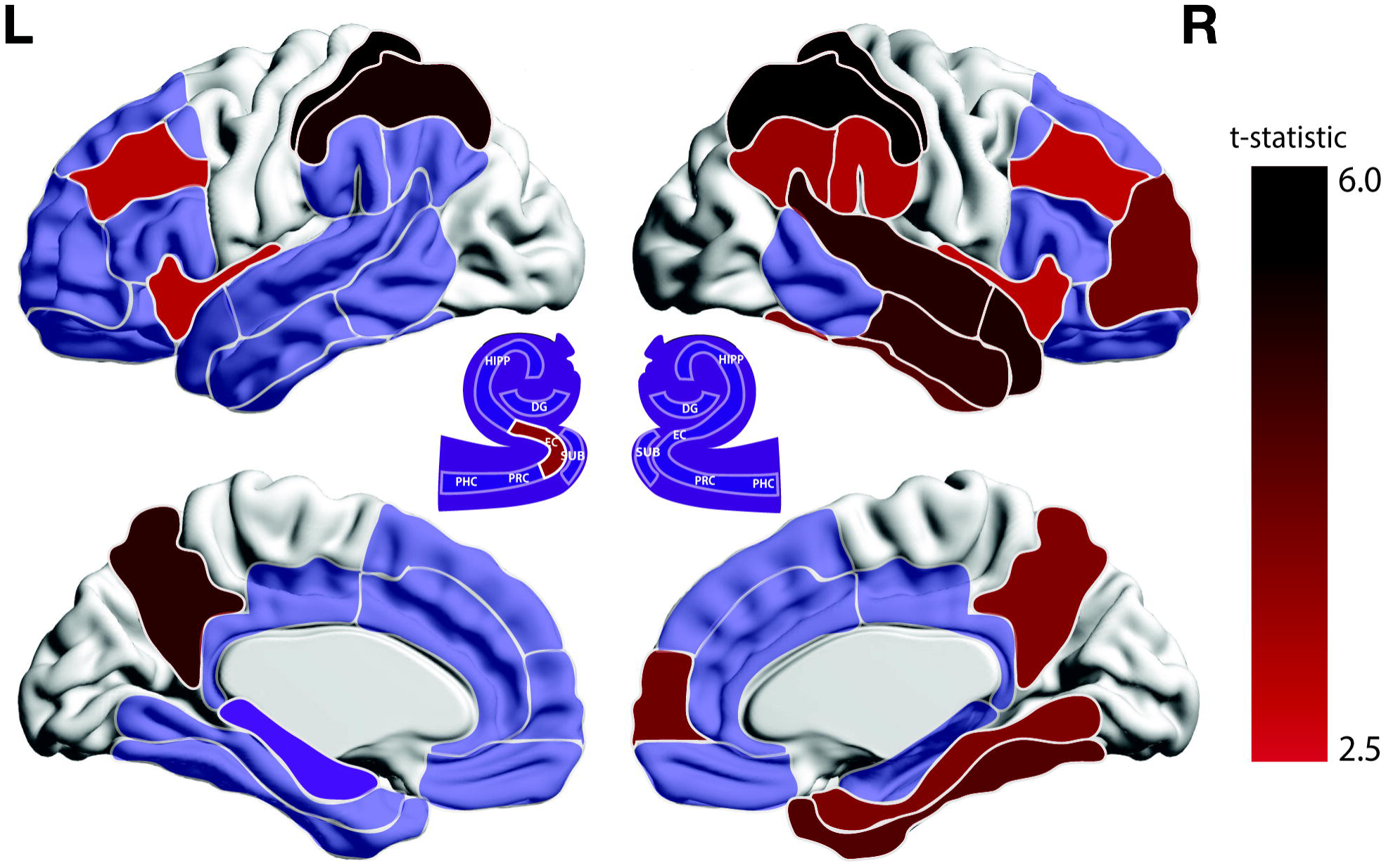
High-Gamma power during arithmetic task versus memory task. Regions shaded red represent those demonstrating significantly more HFA power during the arithmetic task than the memory task. The color bar links the color of a region to the t-statistic for the arithmetic versus memory comparison. The illustration in the center of the brain plots is an illustration of the mesial temporal lobe. Only the left entorhinal cortex is colored in red.

### 2.6 Time series analysis

To examine the time course of gamma power activation during arithmetical processing we averaged the Z value from each intra-electrode comparison of arithmetic versus memory processing across electrodes and subjects within a given brain region while preserving the time series information. Peaks in arithmetic-related activation were identified as those time points in which the Z value from the arithmetic versus memory comparison (HFA) exceeded 90% of the maximum value of this metric across the time series. This technique was chosen instead of isolating a single time point of maximum power to avoid mistaking potentially spurious power spikes for the epoch of maximal sustained regional activity. Data were binned in 2 ms steps. For key regions, we also tested distributions of peak times across subjects (each subject contributing 10 time points of greatest HFA) directly against each other with a t-test. Finally, we compared the distributions of arithmetic-related HFA activity directly between the hemispheres at each time sample, allowing us to plot how hemispheric asymmetry evolved across the time series for each brain region.

### 2.7 Hemispheric analysis

We analyzed differences in activity between the hemispheres in two ways. First, we compared hemispheric data from subjects who had electrodes inserted bilaterally with a paired t-test. Additionally, to take advantage of the full breadth of our data set, we compared activity between hemispheres for each region, allowing every subject to contribute data. We present results from both of these. Additionally, using only subjects with electrode coverage in both hemispheres of a given region, we assessed the difference in power between right and left hemispheres for each time point to assess the trend in hemispheric dominance over time.

### 2.8 Selective anatomical analysis

Based upon pre-existing data indicating that there are differences in the magnitude of arithmetic-related activation along the anterior-posterior axis of the IPS and the NFA (Daitch et al. 2016; Shum et al. 2013), we sought to analyze HFA during arithmetic processing continuously along the y-axis for the IPS, ITG, and fusiform gyrus. We utilized the location of each electrode along the y-axis in Talairach space. For this analysis, all electrodes localized to these brain regions were plotted in a normalized brain viewer to ensure the coordinate values accurately reflected the identified anatomical location. We then separately analyzed arithmetic versus memory HFA at different locations within each region.

### 2.9 Gender analysis

We constructed a mixed effects ANOVA model to predict the magnitude of arithmetic-related HFA across all regions identified in the core arithmetical processing network. Gender and hemisphere were considered fixed effects and each subject contributed a single value for each region. We tested for a primary effect of gender and interaction between gender and hemisphere in this model.

## 3. Results

### 3.1 Behavioral results

With the goal of characterizing a specific brain network for arithmetic processing, we analyzed oscillatory activity recorded as subjects performed a verbal memory task as compared to activity recorded during arithmetical addition. Across several years and eight institutions, we collected data from 343 subjects who performed free recall, and an arithmetical distraction task. Of the 343 subjects who completed the experimental task, 33 were excluded due to right hemisphere dominance as determined by standard pre-surgical evaluation (fMRI or Wada test). Descriptions of average arithmetical performance (as measured by response time) and the correlation between arithmetic performance and memory performance are shown in Figures 1C and 1D. Across these remaining 310 subjects, mean arithmetic problems solving time was 6.55 seconds (median time was 5.43 seconds). Response time correlated negatively with the percent of words properly recalled (meaning faster math performance correlated with better memory performance, *rho*^2^ = 0.1158). Mean percent of arithmetic problems answered correctly was 91.17% (standard deviation of 12.35%) while mean percent of words successfully encoded was 25.28% (standard deviation of 10.75%).

### 3.2 Brain-wide oscillatory power

In our first analysis, we directly compared gamma frequency oscillatory activity for successful memory encoding events with successful arithmetical processing across the two hemispheres for 27 brain regions. For this initial analysis, we separately analyzed data from each hemisphere. Within each recording electrode, we compared high gamma oscillatory power (64 – 128 Hz) during successful arithmetical processing to that during successful memory encoding at the subject level. Figure 2 shows the regions for which high gamma power during arithmetical processing was significantly greater than during memory encoding, correcting for multiple comparisons across all regions with a strict (Q = 0.01) threshold. Consistent with existing data (Daitch et al. 2016), the largest math-related HFA effects were observed in the IPS, SPL, and precuneus, which exhibited bilateral arithmetic-related activation. The latter structure, previously lumped together with more lateral dorsal parietal locations, has not previously been described independently as a part of a parietal circuit underlying arithmetical processing. The arithmetic-specific effect in the IPL, including the angular and supramarginal gyri, was significant in the right but not the left hemisphere. Similarly, in the right hemisphere, we observed significant activation in the superior temporal gyrus. Areas likely involved in numerical recognition (the fusiform and inferior temporal gyri) also exhibited significantly greater activation during arithmetical processing versus verbal memory. The distribution of effects in these brain regions are described below in a more detailed anatomical analysis (Figure 3). The brain regions not previously identified via electrophysiological investigation as part of an arithmetical processing network include the dorsolateral prefrontal cortex, frontal pole, temporal pole, insula and entorhinal cortex, although non-invasive data has implicated prefrontal regions and the insula as part of an arithmetic processing network (Simon et al. 2002). We observed peri-sylvian activation preferentially in the right hemisphere, including the superior and middle temporal gyrus will as the IPL. Activation of mesial temporal structures was specific for the entorhinal cortex; hippocampal regions and parahippocampal cortex did not show an arithmetic-related effect.

**Figure 3:**
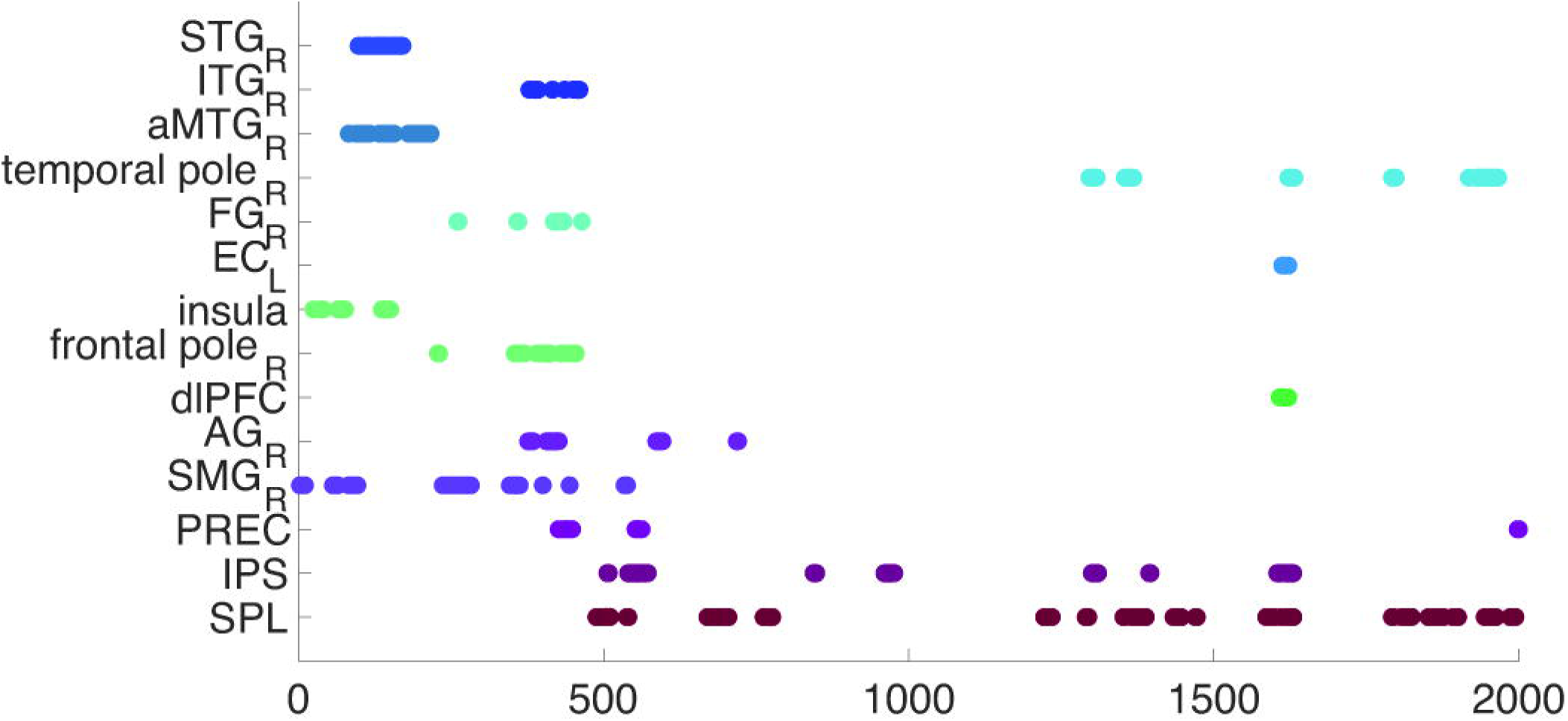
Subregion analysis of the IPS, FG, and ITG. Regions in which we anticipated possible differences along the longitudinal axis are included. The x-axis of each plot is the Talairach x domain. As such, the right of the plots is the right hemisphere, the left of the plot is the left hemisphere. The y-axis of each plot is the Talairach y domain. The color within each plot in each plot represents the fraction of electrodes within the given hemisphere and y-coordinate range that show arithmetic-preferential activity as demonstrated by a z-score > 0. Red boxes represent subregions with more arithmetic-preferential electrodes. Blue boxes represent subregions with more memory-preferential electrodes. Green boxes represent subregions with neither arithmetic nor memory preference.

### 3.3 Spatial analysis

For our next analysis, we wanted to examine more precise spatial activation patterns for a subset of these core arithmetic regions (Figure 3). This was motivated by hypothesized differences in activation along the y-axis for the IPS, ITG, and FG (Daitch et al. 2016; Shum et al. 2013). We began by dividing electrodes located within each region into left and right groups and then divided them up along the y-axis location of each in Talairach space. As each resulting y-axis location had potentially a different number of electrodes contributing, we tested for significance within each subregion by quantifying the percentage of electrodes exhibiting preferential arithmetic related activation (*Z-score* > 0, t-test between arithmetic and memory high gamma power distributions), testing the resulting number against the chance rate using a binomial test. We observed strong activation along the length of the IPS except for the most posterior section on the right (*Talairachy* < −68). In the FG and ITG we observed strong bilateral posterior activation consistent with a “number form area,” although we also observed activation more anteriorly towards the temporal pole (Shum et al. 2013).

### 3.4 Analysis of temporal dynamics

For our next analysis, we sought to assess the timing of activation in each of the brain regions identified as part of the arithmetical processing network (Figure 4). This analysis follows previously published human electrophysiological data (Daitch et al. 2016), although our results reflect subject level aggregation of HFA activation (each subject contributing a single value per region), with at least 16 subjects contributing to data from each brain region and the IPS/SPL including around 40 subjects per hemisphere. We identified peak HFA activation relative to the onset of each arithmetic problem event by normalizing HFA effect size (Z score from arithmetic versus memory test) across the time series and identifying time points at which activation exceeded 90% of the maximum value. Results are shown in Figure 4. Activation peaked in the FG and ITG significantly before the IPS (*p < 0.001*, t-test comparing distribution of peak times across subjects) and the SPL (*p < 0.001*). Entorhinal cortex peak activation significantly preceded FG (*p < 0.001*), IPS (*p = 0.0084*), and SPL (*p < 0.001*) but not ITG. IPS peak activation significantly preceded that of the SPL (*p < 0.001*). Finally, the EC peak significantly proceeded that of the temporal pole (*p = 0.002*) but the timing of the temporal pole peak did not significantly precede that of the SPL (*p = 0.328*).

**Figure 4:**
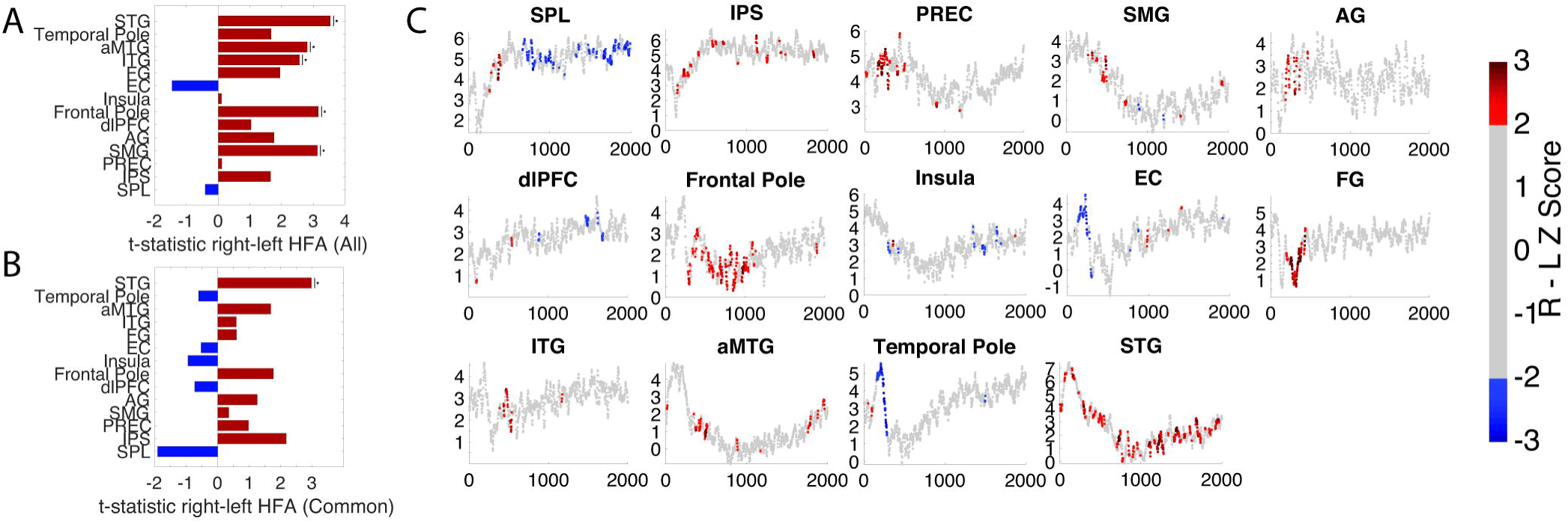
Time of peak HFA for each arithmetic-significant region. Each data point represents a time at which the arithmetic-related high-gamma power of a region is within 90 percent of its maximum across the time series. If the right and left hemisphere of a region demonstrated significantly more activity during arithmetic than memory, the data from each hemisphere were combined prior to the analysis. Otherwise, only the significant hemisphere was analyzed.

### 3.5 Hemisphere analysis

For our next analysis, we sought to directly compare activation in the right and left cerebral hemispheres for the core arithmetical processing regions identified above. We did this in two phases. In the first, we included only subjects in whom electrodes had been inserted into the region of interest in *both* hemispheres; while such data were previously quite rare outside of the temporal lobe, the inclusion of a large fraction of stereo EEG subjects in our dataset afforded us the opportunity to conduct such an analysis. The advantage in this analysis is that possible aberrations in hemispheric specialization due to the underlying clinical situation (epilepsy) would be less of a concern for a paired analysis. However, we also wanted to take advantage of the full richness of our dataset and so we also included all available subjects contributing electrodes to a given brain region into a separate direct comparison of hemispheric activity; subjects were allowed to contribute to both hemispheres if electrodes were represented in both. For the latter analysis, activation was significantly greater for several regions within the right hemisphere, especially for peri-sylvian and temporal regions (FDR corrected *p* < 0.05, paired t–test at subject level). However, only the STG exhibited significantly greater activity in the right hemisphere across both analytical methods (Figure 5A, 5B).

**Figure 5:**
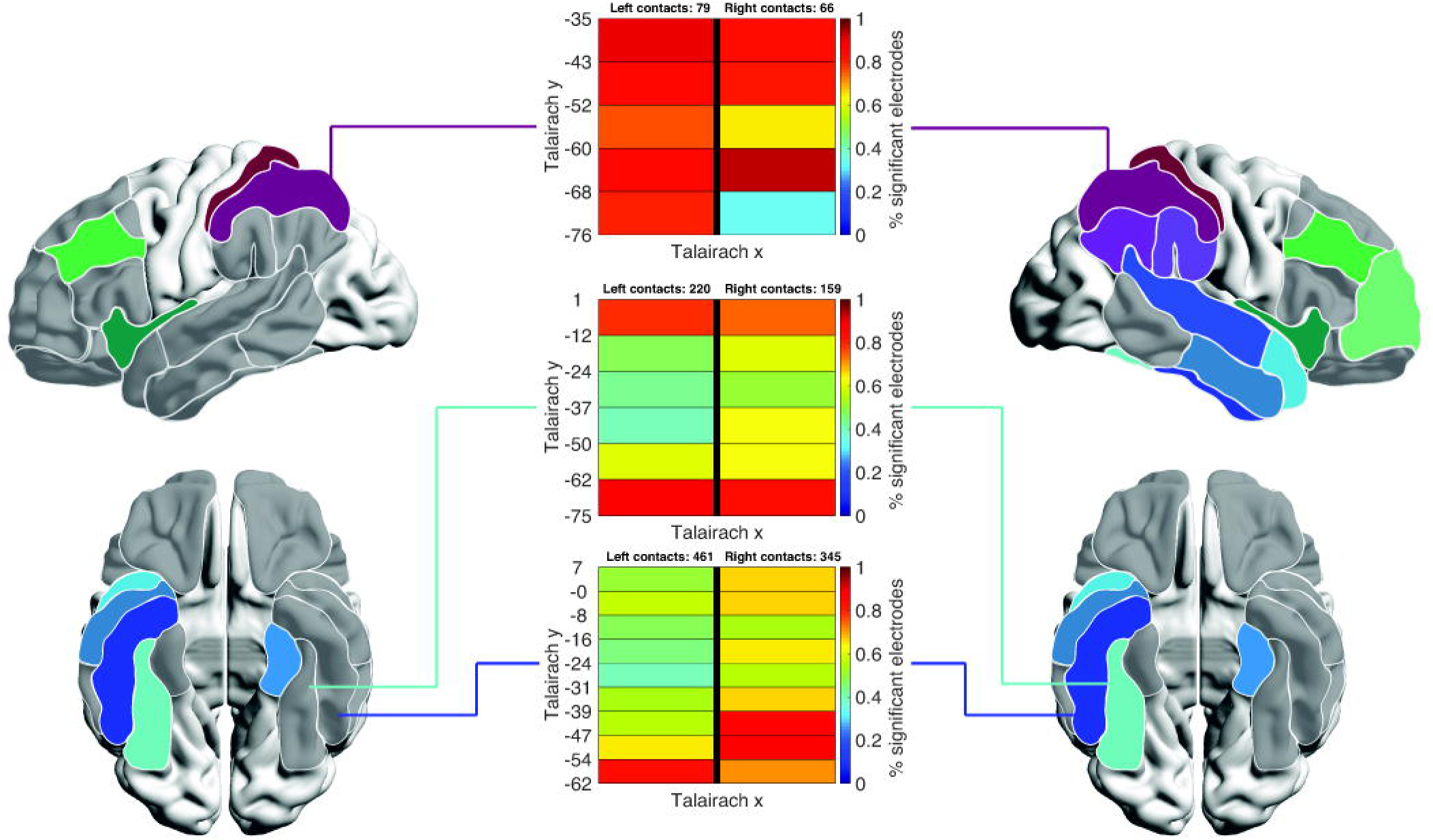
Hemispheric dynamics of each arithmetic-significant region. **A.** Unpaired comparison of right and left hemisphere HFA (all subjects included). **B.** Paired comparison of right and left hemisphere HFA (only subjects with contacts in both hemispheres included). **C.** Plot of arithmetic versus memory power (y-axis, units of z-score) across time (x axis, units of ms) with hemipsheric dominance at each time point represented by the color of that point (units of z-score comparing right to left-sided effects). Red colors represent right-hemispheric dominance as measured by a Z score ≥ 2. Blue colors represent left-hemispheric dominance. Subjects with neurologically determined right hemisphere language dominance were excluded from this analysis.

We next repeated our detailed look at the temporal dynamics of activation within the arithmetic network using a three-dimensional plot in which hemispheric differences (paired t-test between hemispheres, across subjects) are superimposed on the time course of activation within each region (comparison of normalized HFA between arithmetic and memory condition). For these plots, only subjects with bilateral representation for each region are included (see Table 1 for subject counts); statistical comparisons are between successful arithmetic versus memory events (y-axis) and then between the magnitude of these effects in each hemisphere (color axis, Figure 5C). In the SPL, we observed an unexpected transition from predominant right hemispheric activation in the early portion of the timeseries followed by strong left-sided dominance (Z score ≥ 2) beginning 600 ms after stimulus onset. This is in contrast to the IPS, in which a slight right hemispheric predominance persisted across the time-series. Early right-sided predominance followed by bilateral activation was observed for the fusiform and inferior temporal regions, consistent with previous reports (Shum et al. 2013). However, the superior temporal gyrus exhibited predominantly right-sided activation throughout the time series. Both the entorhinal cortex and temporal pole experienced a left-dominant power increase early in the time series.

**Table 1.**
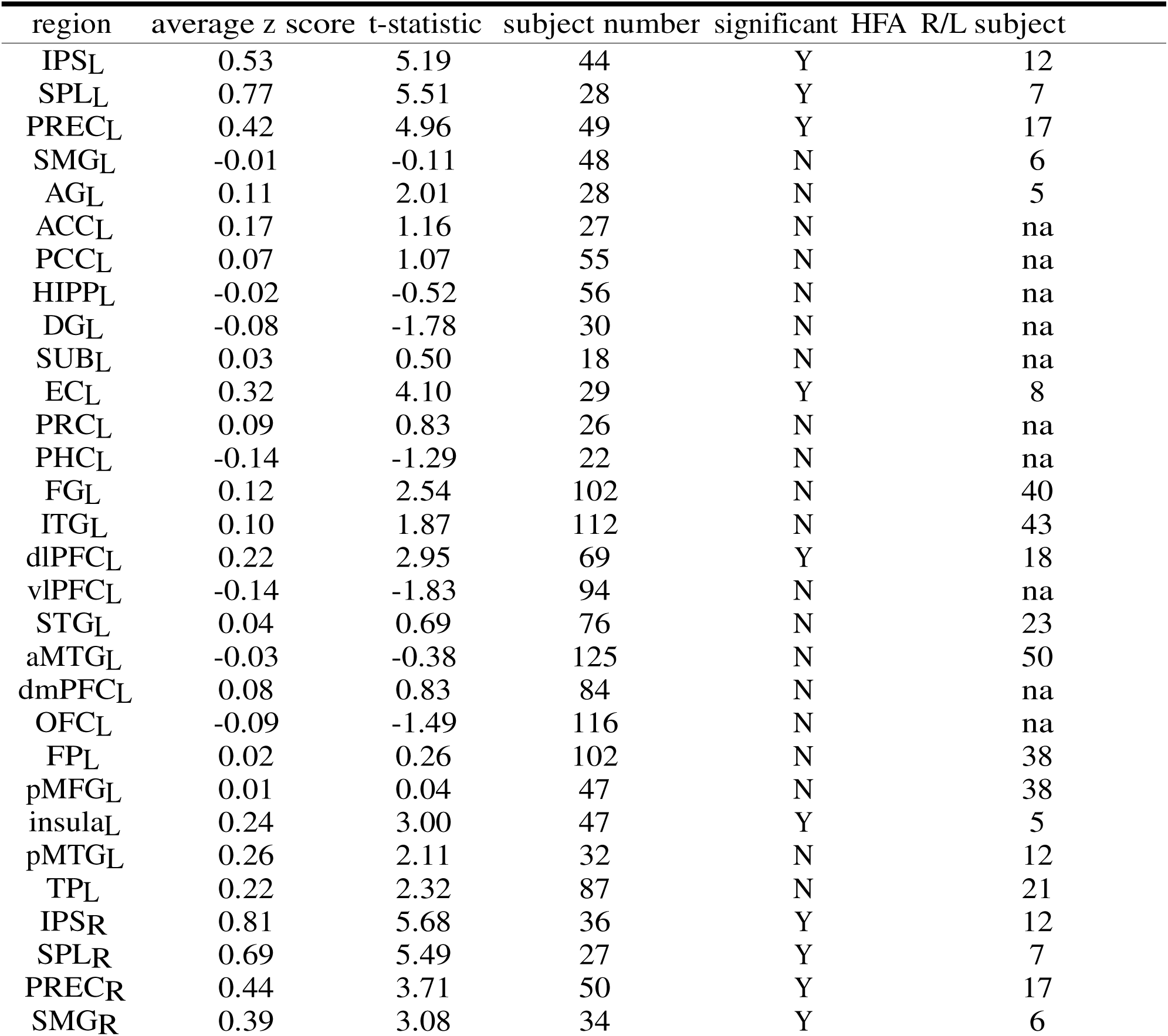
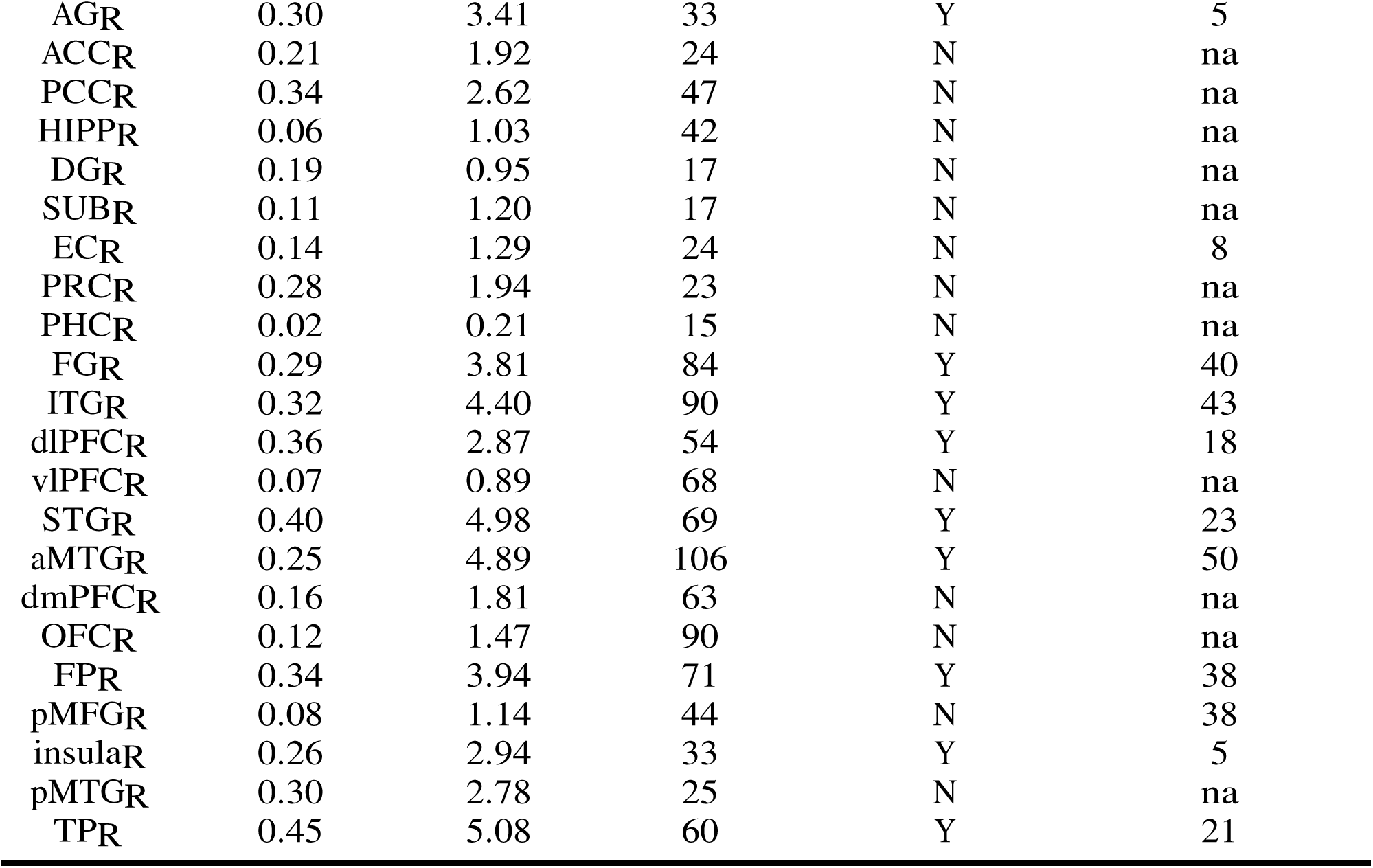

### 3.6 Gender and problem-solving strategy

Our large data set allowed us to look for gender-related differences in activation within the core processing network for arithmetic we describe. This analysis was motivated by an fMRI investigation reporting greater arithmetic-related activation in males in the IPS and angular gyrus (Keller and Menon 2009). Within each region, we constructed a mixed effects ANOVA model, looking for a primary effect of gender and an interaction between gender and hemisphere for predicting the HFA arithmetic-related effect. We saw no significant primary effects of gender (FDR corrected *p* > 0.05), or a gender-hemisphere interaction across our large patient sample (all *p* > 0.1294). Our analysis yields no further support to the notion that problem solving strategy varies between genders.

### 3.7 Correlation of regional HFA effects with behavioral performance

With a large number of subjects contributing data to each region, we sought to quantify the relationship between overall arithmetic performance (as quantified by response time for correct answers) and arithmetic-related HFA activation (Figure 6). We correlated these values across subjects (single value for each region per subject) using a Spearman correlation. Outliers (arithmetic response times greater than 3 median absolute deviations away from the median) were excluded. As expected, the largest correlation values were observed in parietal regions, including the IPS and SPL. However, consistent with differences observed in the hemisphere analysis (Figure 5), we observed a dissociation in results for the left versus right SPL (ρ^2^ = 0.0689 for the left SPL versus 0.0004 for the right SPL). While the SMG had the second greatest correlation with performance (ρ^2^ = 0.0434), this correlation was negative, indicating that individuals with greater activity in this area answered math problems more slowly. Interestingly, the EC and temporal pole, regions implicated in arithmetic reasoning for the first time in our study, demonstrated strong positive correlations with performance (ρ^2^ = 0.0183 and ρ^2^ = 0.0400 respectively).

**Figure 6:**
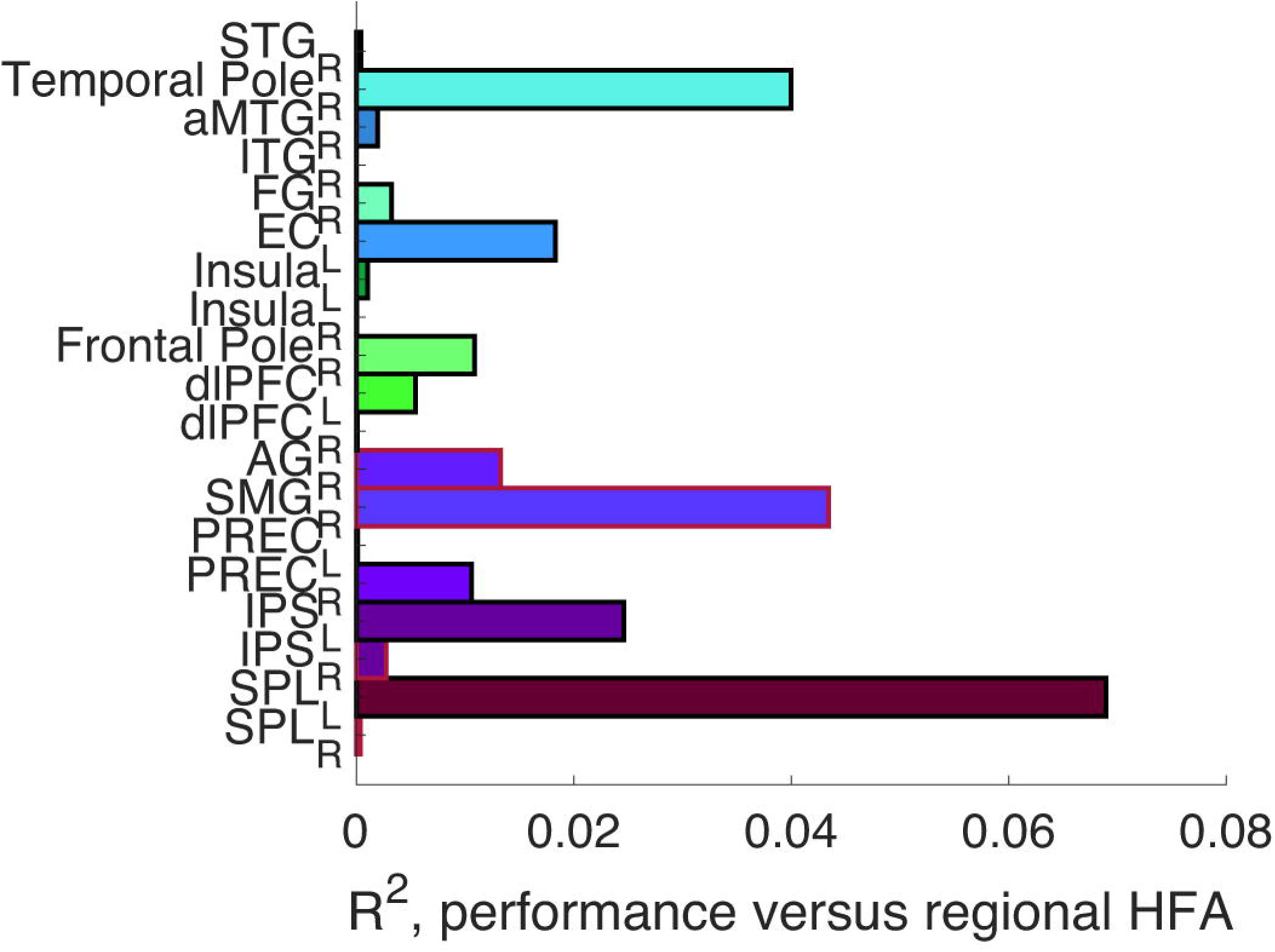
Functional contribution of each arithmetic-significant region. Plot demonstrating the fraction of arithmetic response time that can be explained by the HFA of each arithmetic-significant region. Purple tones represent parietal regions. Green tones represent frontal lobe regions. Blue tones represent temporal lobe regions. If a bar is outlined in red, that region’s HFA correlates negatively with behavioral performance (slower arithmetic response times). Otherwise, increased HFA for a region correlates with faster arithmetic response times.

## 4. Discussion

We describe a human brain network for arithmetical processing based upon aggregated intracranial recordings across 310 human participants. The scope of our data set allowed us to analyze activity across the entire brain, compare activity between the hemispheres, and more precisely define temporal and spatial dynamics within a core arithmetic processing network. We defined this network by identifying brain locations that exhibited significantly greater activation during arithmetic processing as compared to successful memory encoding. Our methods identified regions that were expected a priori to exhibit strong arithmetical activation (IPS and SPL) based upon existing human data and theoretical models of arithmetical processing, but we also identified novel regions (namely the EC, DLPFC, STG) not previously examined using direct brain recordings. The characterization of temporal processing patterns across this network allows us to generate hypotheses about the functional contributions of each region towards arithmetical processing. We also provide the first within-subjects comparison of hemispheric differences across these brain regions. We discuss how our findings inform existing theories of arithmetical processing and examine how these theories may be adapted to incorporate the novel brain areas we identify. It should be noted that evidence exists that higher forms of mathematics such as algebra, topology, and geometry recruit the same basic brain circuit as simple arithmetic (Amalric and Dehaene 2016), motivating the usefulness of using an arithmetic task to probe the electrophysiology of arithmetic specifically and mathematical circuitry more generally.

### 4.1 Triple Code Theory and Three Parietal Circuits

The most influential work in this domain, synthesizing noninvasive and neuropsychological data regarding mathematical processing, is the triple code theory articulated by Dehaene (Dehaene 1992). This theory postulates that arithmetical reasoning first requires translation of arithmetical tasks into a numerical code by which the symbols representing numerical quantities are identified from within the visual stream. From there, depending upon the exact nature of the task, the stimulus may be coded as a semantic fact retrieved directly from memory (akin to word meanings, a verbal code) or may preferentially be translated into a “magnitude code” along an internal mental number line. As with previous findings, our data partially support triple code theory in that we identify distinct areas of activation early in the time series in visual processing areas consistent with the existence of a “number form area” that would be responsible for the initial recognition of the visual input and translation into an Arabic numeral. We observed mixed data regarding the hemispheric preference of activation in these regions. Across all subjects in the ITG, arithmetic related activation was stronger in the right hemisphere, but in the paired analysis (subjects contributing electrodes from both hemispheres simultaneously), no significant differences were observed (Figure 5A, 5B). Previous human electrophysiology experiments have suggested a primarily right hemisphere location for the number form area (Shum et al. 2013), yet subtle differences in the contributions of each hemisphere towards early numerical processing remain an open area for investigation. We did not observe any evidence of expected semantic contributions in the IPL in the left hemisphere. In fact, we observed greater activation in the right hemisphere in this region. Strictly speaking, this is not consistent with triple code theory. However, the use of a verbal memory task as a baseline condition in our experiment likely contributed to this observation; our paradigm is not tuned to detect contributions from semantic language networks towards math processing. Even with this caveat, our observation that the *right* IPL and STG participate in arithmetic processing networks represents new information to integrate into models of how the brain performs math calculations, consistent with recent reports using non-invasive imaging (Amalric and Dehaene 2019). The strongest activation we observed (Figure 2) was in the IPS, SPL, and precuneus, and these regions predict arithmetic performance strongly compared to other regions (Figure 6). We observed divergence in the activation pattern of the IPS, SPL, and precuneus in terms of temporal dynamics and preferred hemisphere. These data suggest that these regions serve different functional roles during arithmetical processing, with the earlier activation of the precuneus consistent with a spatial attention role, which is hypothesized in the three parietal circuits model. However, the SPL and the IPS exhibit more persistent activation throughout the remainder of the time series.

Hemispheric activity differences in the pattern of HFA in these regions suggests divergent roles during arithmetical problem solving. Regarding the IPS, our data generally support bihemispheric activation (Figure 5C), consistent with existing data (Dehaene et al. 2003) and the finding that magnitude information is accessible to both hemispheres in split-brain patients (Cohen and Dehaene 1996).

The SPL, by contrast, exhibited significant hemispheric difference in both gamma activation and in prediction of memory performance (Figure 5C, 6). This implies the SPL complements rather than mirrors the function of the IPS in arithmetic processing. Therefore, while the SPL may participate in quantity manipulation directly along with the IPS, it could also provide a continuous spatial attention signal. This is reconcilable with the SPL’s known role in attention orienting (Corbetta and Shulman 2002) and with the observation that numerosity-specific neurons have not been identified in the SPL unlike in the IPS and frontal cortex (Nieder 2012; Piazza et al. 2004). This may be consistent with a model by which the SPL in the dominant hemisphere is actively *suppressing* competing (external) sensory inputs to favor attention to internal numerical representations during quantity manipulation (Spreng et al. 2013), a proposal which may be consistent with the specific contributions of this region to working memory (Koenigs et al. 2009), while the role of specifically, *numerically-relevant* attention may be served by the precuneus, which in our data exhibited bilateral activity.

The competing view is that the SPL directs attention towards the internal numerical representation (rather than away from something else) (Dehaene et al. 2003). This suggestion stems the discovery of overlap in the cortex of numerical comparison and spatial attention functions (Pinel et al. 2001) and from a lesion study that demonstrates SPL damage leads to ipsilateral deviation on numerical bisection tasks (Zorzi et al. 2002). These findings predict that both the left and right SPL should be activated in any cognitive task requiring the internal number line in order to attend to the full domain of values. However, our data show preferential activity of the left SPL, which would translate into preferential attention to the right side of the internal number line representing greater numerosities. This could mean that our data favor the theory that the left SPL inhibits redirection of attention away from the internal number line. However, our observations may point to a common mental strategy among participants to draw internal number lines with as small of a range at possible. Since we asked our subjects to solve a two-step question, the number line drawn for the first step of the problem may not have included the value of the eventual answer following the second step. This would require careful attention to the upper bound of the first number line and careful drawing of the upper bound of the second number line. Further characterization of these subtle hemispheric differences using alternate mathematical tasks will be of strong interest.

### 4.2 IPL during arithmetical processing

Dehaene proposes a dorsal versus ventral distinction in parietal contribution to arithmetical processing with more ventral parietal areas (angular gyrus) being activated during mathematical fact retrieval-based problem-solving (Dehaene et al. 2003). He hypothesizes that this (semantic) circuit is preferentially activated in the dominant hemisphere. This is in contrast to more dorsal regions, namely the IPS, which exhibits bilateral activation for mathematical problems that require numerical coding or estimation. Our observations diverged from this model most strongly in terms of the angular gyrus and semantic networks more generally. We did observe arithmetic related activation in the *right* angular gyrus, and the timing of this activation (400 ms after item presentation) is consistent with what one may predict based upon its hypothesized role within the Dehaene model. As discussed above, this may be attributable to our comparison condition which was a verbal memory task: contributions of the left angular gyrus to verbal memory and arithmetical processing may be similar. Also, the multistep nature of our arithmetical task may have favored magnitude based arithmetical reasoning rather than fact retrieval using semantic networks. However, another possible interpretation is that a semantic network for mathematical fact retrieval relies on bilateral cortical areas more strongly than spoken language, reflecting the necessary interaction between symbolic meanings in the numerals and the spatial processing regions necessary for many strategies for solving arithmetic problems. Another possibility is that these regions represent contributions from spatial processing areas without any specific semantic role. Further investigation of contributions of this region to geometrical tasks or with a more precise comparison with straightforward fact retrieval will be needed to elucidate this finding using electrophysiological data.

### 4.3 Mesial temporal circuits during arithmetical processing

The strong IPS/SPL and ITG/FG activation and the temporal activation patterns we observed in these regions is highly consistent with both invasive and noninvasive existing human literature (Amalric and Dehaene 2016; Daitch et al. 2016), indicating that our methods were able to identify brain regions for which a strong a priori assumption existed for participation in arithmetic processing networks. And yet, we also observed strong activation in the left entorhinal cortex, although not in other mesial temporal regions. We hypothesize numerous possible roles for the EC. One is that it participates in a fronto-temporal working memory system involved in maintaining externally directed attention and manipulation of intermediate representations for the arithmetic task. There are conflicting data in the existing literature in this regard, with single unit electrophysiological evidence of participation in a working memory buffer system for the mesial temporal lobe (Kornbilth et al. 2017) but also observations that MTL lesions do not lead to working memory deficits (Cave and Squire 1992).

Another possible explanation for our observation is that the EC is involved in translating the Arabic numerical code processed initially in the FG/ITG into a spatial code compatible with the spatial processing regions of the precuneus, SPL, and possibly of the IPS itself. Clearly, there is long-standing evidence that the entorhinal cortex is activated during both real and imagined navigation (Hafting et al. 2005; Jacobs et al. 2013). It may be the case that (depending upon an individual’s arithmetic processing strategy) numerical processing utilizes some of the same underlying circuitry. Evidence in favor of this latter hypothesis we believe comes from the timing of activation of the EC in the left hemisphere (Figure 5C), concurrently with early activation of the ITG (p-value for test for peak activation time of ITG versus EC = 0.131) but before activation in the SPL (*p < 0.001*). We observed bilateral EC activation later in the time series (around 1600 ms, Figure 4) that was coincident with times of peak activation of the temporal pole and the DLPFC. This observation is more consistent with a working memory-based hypothesis (Egorov et al. 2002). Directly comparing activation in arithmetical processing versus spatial navigation or spatial object memory on a within-subjects basis may prove insightful for elucidating this question.

Additionally, both the medial and lateral entorhinal cortex have demonstrated function that could support arithmetic processing. The properties of the grid code of hexadirectional cells in the dorsolateral medial entorhinal cortex (dMEC) are similar to those of the fixed-base numeral systems we use in mathematics (Fiete et al. 2008). Therefore, perhaps the dMEC is able to explicitly represent numerosity in addition to position in space. The lateral entorhinal cortex was recently implicated in inherently representing time through the encoding of experiences (Tsao et al. 2018). Since performing arithmetic inherently requires completion of a sequence of operations, it is possible the lateral entorhinal cortex ensures that operations are performed in the proper order.

### 4.4 Frontal lobe in arithmetical processing

The DLPFC plays a role in retention of information in working memory (Curtis and D’Esposito 2003). It is thought that this is achieved by inward attention to semantic and/or sensorimotor representations, though the exact neuronal mechanism of this is a matter of current debate (D’Esposito and Postle 2015). Activation in the DLPFC for arithmetical processing has not previously been reported using direct electrophysiological recordings in humans but was not an unexpected finding based upon the nature of our task (multistep addition) which presumably involves strong activation of working memory circuits to maintain intermediate values for further manipulation in mathematical regions. Further, the timing of activation of the DLPFC favors a role in working memory after initial stages of arithmetical processing in the IPS and SPL (Figure 4). On the other hand, primate data exist to support direct numerical processing within the DLPFC, as numerically sensitive neurons have been identified in this brain area (Nieder 2012). The timing of activation that we observed could also be consistent with a role for the DLPFC in more challenging arithmetical operations that recruit extra–parietal resources along with the IPS. The frontal pole has been implicated as the site of ‘cognitive branching’ (Koechlin and Hyafil 2007). According to this model, the frontal pole allows one to place a current task set in a pending state in order to address a more immediate concern with its own task set. In our experimental task, such a function may be necessary due to the multi-step additional problems the subjects performed, or because of the context of the arithmetic task within a verbal memory paradigm (necessitating switching from one task to another).

### 4.5 Spatial distribution of arithmetic-related activity in the inferior temporal lobe

Our data set allowed us to investigate the spatial distribution of numerical activation in the inferior temporal lobe, including hemispheric asymmetries. We observed that for both mesial and lateral areas (fusiform gyrus versus ITG), there was arithmetic related activation anteriorly (within 5 cm of the temporal pole) as well as posteriorly (in a putative number form area) as expected. Previous authors examining intracranial data describe the phenomenon by which numerically relevant HFA increases are highly restricted in space in this brain region (Shum et al. 2013). This may be true at the level of individual subjects; however, our data indicate that in the aggregate arithmetic selective regions are visible across a broad number of individuals, and further that this mathematically sensitive area extends anteriorly. This observation has implications for expected deficits following standard temporal lobe surgery, as a typical temporal lobectomy operation will extend to this area in the nondominant hemisphere. It implies that numerical processing tasks could be included in functional mapping routines, both intraoperatively and extra operatively.

### 4.6 Other temporal lobe structures during arithmetical processing

In addition to the expected activation in the ITG/FG, several other temporal lobe structures showed significant arithmetic-related activity. These regions included the right STG, right anterior MTG, and right temporal pole (TP). The superior temporal gyrus activation could be explained in several different ways. Firstly, the right anterior STG is implicated in solving problems with insight, colloquially known as an ‘Aha!’ moment (Jung-Beeman et al. 2004). However, if this were the function of the STG, we would expect the activation peak to be much later in the time series, not well before the brain regions involved in magnitude code manipulations (IPS, SPL, precuneus) demonstrate sustained activity (Figure 4). Conversely, the STG effect could be related to its role in stimulus driven attentional control. Disturbance of the right STG with TMS have been shown to reduce performance in a complex visual search task (Ellison et al. 2004). The right temporo-parietal junction, which includes the posterior STG and the IPL, is implicated in detecting unattended or infrequently encountered stimuli independent of their modality or spatial orientation (Corbetta and Shulman 2002), which may be related to our paradigm in which arithmetic problems were presented within the greater context of a memory task. This also provides an additional explanation for the right SMG/AG activation previously discussed. A role in attention orientation could explain its activation early in the time series (Figure 4, 5C). Finally, STG activation can be explained by its role in language perception. The left STG serves as a substrate for auditory short-term memory and written sentence comprehension (Leff et al. 2009). It also allows for mental speech rehearsal (Buschsbaum et al. 2001). Since the comparison task in our analysis was a verbal memory task, it is possible that significantly more activation in the right STG during arithmetical problem solving signifies leftward lateralization of STG activity during this comparison task.

As compared to more posterior temporal structures, existing anatomical data more strongly support a role for the temporal pole (TP) in numerical processing, consistent with our findings. Diffusion tensor imaging tractography results have revealed two distinct connections of the temporal pole to other major brain regions via the middle longitudinal fasciculus (MLF). The first runs ventrolaterally from the STG and TP to the angular gyrus while the other runs dorsomedially and connects the STG and TP to the superior parietal lobule (Makris et al. 2013). Early activation in the temporal pole relative to the IPS, SPL, and precuneus (Figure 5C) may support a hypothesis that this region (along with the co-activating entorhinal cortex) directs the SPL to the proper location along the mental number line mediated by MLF connections. Connections to the angular gyrus in turn would support translating the numerical symbols into a semantic code consistent with the Dehaene model (Dehaene et al. 2003).

### 4.7 The insula during arithmetical processing

There exists substantial noninvasive evidence of mathematics-related insular activation. It is active during subtraction (Simon et al. 2002). It also demonstrates strong distance-dependent activity recovery during presentation of numerical deviants in adults (Piazza et al. 2007). This means that after adapting to repeated stimuli of the same quantity, upon presentation with a different quantity, the insula activates proportionally to the distance between the numerosity to which it is adapted and the new numerosity. This metric is used to identify regions involved in abstract encoding of numerical magnitude. In Piazza et al. 2007, the insula showed a stronger effect than the parietal cortex. In children aged 4-11, the bilateral insula activated preferentially to numbers compared to faces, shapes and words, demonstrating that the insula’s role in mathematical processing begins early in development (Emerson and Cantlon 2011). However, this region also activates when experiment participants anxiously anticipate being asked to perform math problems. When anticipating the need to perform a math task, but not during math performance itself, individuals with an aversion demonstrate insular activation which may be related to a subjective experience of pain (Lyons and Beilock 2012). In our study, the insula activated earlier than the FG and the ITG (Figure 4 and 5C), regions classically involved in interpreting Arabic numerals, presumably among the first steps of arithmetic processing. It is possible this early activation is a fast recognition of the magnitude represented by the Arabic numerals, but it is perphaps more likely that this is a signature of anxious math anticipation. Activation was bilateral, consistent with the aforementioned studies (Figure 5C).

### 4.8 Regional influence on arithmetic performance

Figure 6 illustrates how the HFA in each region contributes to subject level variance in the speed of arithmetic performance. By lobe, the parietal lobe had the strongest correlation with arithmetic speed followed by the temporal lobe. The frontal lobe demonstrated very little contribution to performance variance. Interestingly, while left SPL and left IPS HFA correlated positively with arithmetic performance, their right hemisphere counterparts correlated negatively. The SPL result is consistent with left hemisphere dominant HFA in the SPL during the arithmetic task (Figure 5C). It was surprising to observe significant association with performance in only the left SPL. We propose this suggests that multi-step addition predominantly favors representations on the upper end (right side) of the internal number line, an abstraction generated by the IPS but attended to by the left SPL. Additionally, the temporal pole and entorhinal cortex both demonstrated strong positive correlations with arithmetic performance. This highlights the importance of further investigating the proposed EC-temporal pole-SPL activation axis previously discussed.

## 5. Conclusion

Using an unparalleled data set of 310 subjects with intracranial electrodes performing a multistep addition task, we characterize the spatial and temporal properties of an arithmetic processing network. Our methods emphasize the contributions of the IPS and SPL consistent with previous observations, but we describe previously unreported arithmetic-related activity in the nondominant temporal lobe, the entorhinal cortex, and the DLPFC. We also report hemispheric differences in SPL activation. Our findings support core hypotheses of triple code theory, but also suggest how this theory may be modified or combined with other characterizations of parietal lobe activity during cognitive processing.

## 6. Acknowledgements

This work was supported by the UTSW/THR Clinical Scholar Program and the DARPA Restoring Active Memory (RAM) program (Cooperative Agreement N66001-14-2-4032). The views, opinions, and/or findings contained in this material are those of the authors and should not be interpreted as representing the official views or policies of the Department of Defense or the U.S. Government. The authors declare no competing conflicts of interest.

